# Mechanical forces pattern endocardial Notch activation via mTORC2-PKC pathway

**DOI:** 10.1101/2024.04.01.587562

**Authors:** Yunfei Mu, Shijia Hu, Xiangyang Liu, Xin Tang, Jiayi Lin, Hongjun Shi

**Affiliations:** Fudan University, Shanghai, 200433, China; Key Laboratory of Growth Regulation and Translational Research of Zhejiang Province, School of Life Sciences, Westlake University, Hangzhou 310024, China; Westlake Laboratory of Life Sciences and Biomedicine, Hangzhou, Zhejiang, China; Institute of Basic Medical Sciences, Westlake Institute for Advanced Study, Hangzhou, Zhejiang, China

**Keywords:** Cardiac patterning, Endocardium, EMT, Notch, Shear stress, Mechanosensing

## Abstract

Notch signaling has been identified as a key regulatory pathway in patterning the endocardium through activation of endothelial-to-mesenchymal transition (EMT) in the atrioventricular canal (AVC) and proximal outflow tract (OFT) region. However, the precise mechanism underlying Notch activation remains elusive. By transiently blocking the heartbeat of E9.5 mouse embryos, we found that Notch activation in the arterial endothelium was dependent on its ligand Dll4, whereas the reduced expression of Dll4 in the endocardium led to a ligand-depleted field, enabling Notch to be specifically activated in AVC and OFT by regional increased shear stress. The strong shear stress altered the membrane lipid microdomain structure of endocardial cells, which activated mTORC2 and PKC and promoted Notch1 cleavage even in the absence of strong ligand stimulation. These findings highlight the role of mechanical forces as a primary cue for endocardial patterning and provide insights into the mechanisms underlying congenital heart diseases of endocardial origin.

## 1. INTRODUCTION

The formation of the mammalian heart begins with a simple linear heart tube which undergoes complex morphogenic processes to ultimately reach the adult four-chamber configuration. In mice, the pattern of the adult heart becomes apparent at embryonic day (E) 9.5. At this stage, increased cell proliferation and sarcomeric development at the outer curvature of the primary heart tube lead to the formation of working chamber myocardium, while less proliferative and less differentiated myocardium at the inner curvature characterizes the outflow tract (OFT) and atrioventricular canal (AVC) ^1, 2^. Between E9.5 and E10.5, endocardial cells in the OFT and AVC regions undergo extensive endothelial-to-mesenchymal transition (EMT), forming endocardial cushions. Further growth and remodeling of these endocardial cushions contribute to the septation of the OFT and cardiac chambers and give rise to cardiac valves ^3^. Malformation of the endocardial cushions underlies the mechanism of the majority of human congenital heart defects (CHD).

Multiple interactive signal transduction pathways are involved in patterning valve versus chamber myocardium and regulating EMT. At E9.5, expression of BMP2 in the AVC and OFT myocardium ^4, 5^ directly activates TBX2 expression, which, in turn, activates TGFβ2 expression ^6^. Both BMP2 and TGF-β2 enhance Snail1 expression and stimulate EMT ^4, 7^. Notch signaling is also required for EMT. At E9.5, NICD (Notch1 intracellular domain) expression is the highest in endocardial cells of the AVC and OFT region, while in the ventricle, NICD is more restricted to the endocardium at the base of developing trabeculae ^8,9^. Notch, through the transcriptional factor CSL, directly activates Snail2 expression, leading to the repression of VE-cadherin expression ^7, 10^. Genetic abrogation of Notch signaling blocks EMT in the AVC endocardium, whereas ectopic endocardial NICD expression leads to partial EMT of ventricular endocardial cells outside AVC ^10^ ^11^.

Despite the well-characterized role of Notch in endocardial patterning and EMT, little is known about the mechanism that specifically activates Notch in the areas of endocardium that undergo EMT. Among various Notch receptors and ligands, Notch1, Dll4, and Jag1 are expressed in the endocardium at the onset of EMT ^10, 12^. Notch1 transcript appears to be uniformly expressed throughout the endocardium at E9.5, while Dll4 is most concentrated in the ventricular endocardium at the base of the trabeculae ^9^. Jag1 is highly expressed throughout the myocardium, with only a few individual AVC endocardial cells positive for Jag1 ^13, 14^. Conditional deletion of Dll4 in all endothelial cells results in the loss of classic EMT markers and complete absence of AVC cushions at E9.5 ^13^. However, it remains unclear whether this impact on EMT is specifically due to the loss of Dll4-Notch signaling at cushion endocardium or a consequence of global circulatory failure resulting from vascular deformation in Dll4-deficient embryos. Therefore, the currently known expression pattern of Notch receptors and ligands does not fully explain the specific pattern of Notch activation in the endocardium.

Studies in zebrafish embryos have revealed the essential role of various mechanosensitive ion channels in modulating Notch1b receptor expression in the AVC endocardium in response to increased shear stress in this area ^15–17^. However, as mentioned earlier, in the developing mouse endocardium at the onset of EMT, Notch1 is uniformly expressed throughout the endocardium despite restricted NICD in AVC and OFT endocardium. Thus, the equivalent mechanosensitive pathways controlling region-specific pattern of Notch activation, and hence EMT, in the mammalian endocardium are still unclear. To address this question, we transiently blocked the heartbeat of mouse embryos at E9.5 *in vivo* using the class III antiarrhythmic drug dofetilide, which selectively blocks the rapid component of the delayed rectifier K^+^ current (I_Kr_) in cardiomyocytes ^18^. We found that Notch activation in the vascular endothelium is dependent on Dll4, whereas in the endocardium, the overall reduction of Dll4 expression creates a ligand-depleted field that allows establishment of a specific Notch activation pattern in the AVC and OFT regions in response to increased shear stress. Strong shear stress in these valve-forming areas alters the membrane lipid microdomain structure of the endocardial cells, which then activates mTORC2 and PKC, promoting Notch1 cleavage even in the absence of strong ligand stimulation. The results uncovered a new mechanism whereby mechanical force serves as a primary cue for endocardial patterning in mammalian embryonic heart.

## 2. STAR METHODS

### 2.1. Reagents

A list of the reagents used in this study is provided in Table S1.

### 2.2. Animals

All mice used in this study were housed at a constant temperature (23°C) and humidity (∼50%) with a 12-h light/dark cycle and ad libitum access to food and water. All animal experiments were performed in accordance with the protocol approved by the Westlake University Institutional Animal Care and Use Committee (approval # 21-007-SHJ). Noon of the day of vaginal plug detection was defined as embryonic day (E) 0.5. For dofetilide treatment, mice were gavaged at E9.5 with a single dose of dofetilide at 2 mg/kg dissolved in saline. For phorbol 12-myristate 13-acetate (PMA, 2 mg/kg) administration, mice were gavaged at E9.5 with a single dose of PMA dissolved in CMC-Na. Verapamil was given by intraperitoneal injection at E9.5. At the indicated date post-treatment, the dams were sacrificed by cervical dislocation, and the embryos were harvested for further analysis. All sources of our mouse lines are listed in Table S1. Knockout mice generated in-house are described and validated in Figure 3—Figure supplement 2E.

### 2.3. Fetal echocardiography

Ultrasound scanning was performed using the Vevo 300 high-frequency ultrasound machine (FUJI FILM Visual Sonics Inc., Canada). *In utero* echocardiography of fetal mice was conducted at E9.5. Transducer MX-550D had a central frequency of 40 MHz with axial and lateral resolutions both about 30 μm. The pregnant mice were sedated using 2% isoflurane and kept at 1% isoflurane to maintain body temperature at 35.5-37°C and heart rate at 400 to 500 beats per minute (bpm). Pulse-wave Doppler was used to measure velocity and time interval parameters to measure the blood flow velocity and time interval parameters of the fetal hearts. All fetuses were scanned twice in order and their *in utero* positions were noted. For embryos of *cTnT^cre^* x *Tnnt2 ^flox/flox^*, after completing the Doppler ultrasound imaging, the embryos were dissected out with the recorded positions, and embryonic tissues were collected for genotype identification to correlate with the heart rate data. During the measurements, observers were blinded to the assigned groups. Analysis was performed with Vevo Lab 5.7.1.

### 2.4. Fetal heart morphology

Fetal hearts were imaged using the Zeiss Lightsheet Z.1 microscope. Embryos were harvested at E18.5, and hearts were dissected in phosphate-buffered saline (PBS), fixed overnight in a mixed solution of 10% neutral buffered formalin and 2.5% glutaraldehyde, then rinsed twice in PBS, and dehydrated in 50, 75 and 100% ethanol for 30 mins each at room temperature (RT). Samples were then transferred into a specially designed glass tube containing 100 μL of BABB solution (1:2 benzyl alcohol: benzyl benzoate) and incubated for 30 mins to clear the sample. The glass tube was mounted into the sample chamber which was filled with 87% glycerol (RI∼1.45). Hearts were scanned from the apex to the great arteries for tissue autofluorescence using the 561 nm laser line and the detection optics 5x/0.16 (n=1.45). 3D reconstruction of the image stacks and morphological analyses were performed with Imaris 9.3 software.

### 2.5. *Ex vivo* embryo culture

Mouse embryos at E9.5 were carefully dissected in Hanks Buffer containing calcium and magnesium to remove decidua without damaging their yolk sacs and placentas. They were then cultured in 1 mL embryo culture medium (50% rat serum, 50% DMEM), pre-saturated with a gas mix of 95% O_2_ and 5% CO_2_ in a 5 mL volume glass tubes. Tubes were sealed and placed in an rotatory shaker maintained at 37 °C and 50 rpm. For pharmacological treatment, stauroporine (100 nM), wortmannin treatment (2 μM), dofetilide (0.2 ug/ml), or water-soluble cholesterol (1 mg/ml) was diluted in embryo culture medium to the desired concentrations.

### 2.6. Immunofluorescence and *in situ* hybridization

E9.5 and E10.5 wild embryos were dissected in PBS, fixed in 4% paraformaldehyde (PFA) overnight at 4 °C, dehydrated in ethanol series, cleared in xylene, and embedded in paraffin. Tissues were sectioned at 6 μm thickness. Paraffin sections were subjected to a 20 mins heat-induced antigen retrieval using a citric acid solution (pH 6). The sections were blocked with 5% normal donkey serum (NDS) in Tris-buffered saline with 1% Tween-20 (TBST) for 1 hour, followed by overnight incubation at 4°C with primary antibodies diluted in the blocking buffer. Bound primary antibodies were detected using fluorescently conjugated corresponding secondary antibodies. For detection of low-abundance targets such as NICD and phospho-PKC^Ser660^, tyramide signal amplification (TSA) was applied. A list of the antibodies used in this study are provided in Table S1. Confocal microscopy images were captured on a Zeiss LSM 800 and image analysis was performed using Zen 2.3. Fluorescent intensity was calculated using ImageJ software.

To analyze NICD expression in E8.5, E9.5 mouse hearts, whole mouse embryos were fixed in 4% PFA at 4°C overnight. After blocking in 5% NDS, 0.5% Triton X-100 in TBS at 4°C overnight, samples were incubated at 4°C overnight with the primary NICD antibody diluted in blocking buffer (1:400). After washes, embryos were incubated with donkey anti rabbit HRP (1:500) 4°C overnight. Embryos were then washed and reacted with a tyramide-biotin solution at RT for 30 minutes. After that, samples were washed and incubated with Cy^TM^3 streptavidin (1:500). Stained embryos were dehydrated in methanol, cleared in BABB and imaged by Zeiss LSM 800.

Whole mount in situ hybridization of *Dll4* were performed using the HCR probe set according to the manufacturer’s instruction (Molecular Instruments, Los Angeles). Stained embryos were dehydrated in methanol, cleared in BABB and imaged by Zeiss LSM 800.

### 2.7 Scanning electron microscopy

E9.5 embryos are fixed with a solution containing 2% paraformaldehyde and 2.5% glutaraldehyde (pH 7.2) at 4°C overnight. The fixed embryos are then washed three times with 0.1M phosphate buffer (PB, pH 7.2–7.4) at 4°C for 10 minutes each. Subsequently, the samples are further fixed with 1% osmium tetroxide in 0.1M PB on ice for 1 hour. After fixation, they are rinsed three times with double-distilled water (ddH₂O) at room temperature for 15 minutes each. Dehydration is carried out at room temperature using a graded ethanol series: 30% ethanol for 10 minutes, 50% ethanol for 10 minutes, 70% ethanol for 10 minutes, 95% ethanol for 10 minutes, and absolute ethanol three times for 10 minutes each. Then the dehydrated samples are subjected to critical point drying. Samples were attached to a sample holder with double-sided carbon tape, and coated with a 10–15 nm thick metal film for SEM imaging.

### 2.8 Statistics

Difference in expression levels were tested using two-tailed t test. Two-sided Fisher’s exact test was used to compare the heart defect rates between two groups. P value <0.05 was considered significant. All statistical analyses were performed in GraphPad Prism 8 software.

## 3. RESULTS

### 3.1. Notch is strongly activated in the AVC and proximal OFT endocardium in spite of weak ligand expression at the onset of EMT

Although Notch1, Dll4, and Jag1 have been reported as the principal receptor and ligands expressed in the endocardium ^12, 13, 19^, the relationship between the Notch activity and its ligand expression in various parts of the E8.5-9.5 endocardium have not been examined in sufficient details. To investigate whether regional-specific Notch activation is due to differential receptor or ligand expression, we performed whole-mount *in situ* hybridization and immunofluorescence staining of Notch1, NICD, Dll4, and Jag1. Both *Dll4* transcripts and NICD were uniformly positive throughout all endothelial lining of the dorsal aorta and heart at E8.5 (Figure 1—Figure supplement 1A, Video S1, S3). However, at E9.5 when endocardial EMT starts, distinct patterns of NICD and Dll4 expression appeared (Figure 1B and Video S2, S4). Based on the expression pattern of NICD and Dll4, overall, the endocardium at E9.5 can be viewed as three distinct types as summarized in Figure 1A. Type I is Dll4-high and NICD-high, including the endocardium lining the distal OFT, the base of the trabeculae, the dorsal wall of the left atrium, and the right atrium. These endocardial cells normally do not undergo EMT. Type II is Dll4-low and NICD-high, including the AVC and proximal OFT endocardium. Endocardial cells in these regions undergo EMT to form endocardial cushion cells. Type III is Dll4-low and NICD-low, including the endocardium flanking the AVC and on the top of ventricular trabeculae (Figure 1B and Figure 1-Figure supplement 1C). Jag1 was highly expressed in the myocardium but expressed at a very low level in a small subset of the OFT and AVC endocardial cells (Figure 1—Figure supplement 1A), consistent with the previous report ^13^. Vascular endothelium continued to express high levels of Dll4 and NICD uniformly, similar to type I endocardial cells. Dll4 protein staining pattern overlapped with the *Dll4* transcript and also agreed with VEGF receptor-2 (Flk1/KDR) protein expression throughout the cardiovascular endothelium at both E8.5 and E9.5 (Figure 1—Figure supplement 1B), consistent with the regulatory role of VEGF signaling in Dll4 expression ^20^. Thus, the Dll4 and NICD expression patterns were disconnected in the type II endocardium at E9.5, suggesting the existence of additional mechanisms for Notch activation.

**Figure 1:**
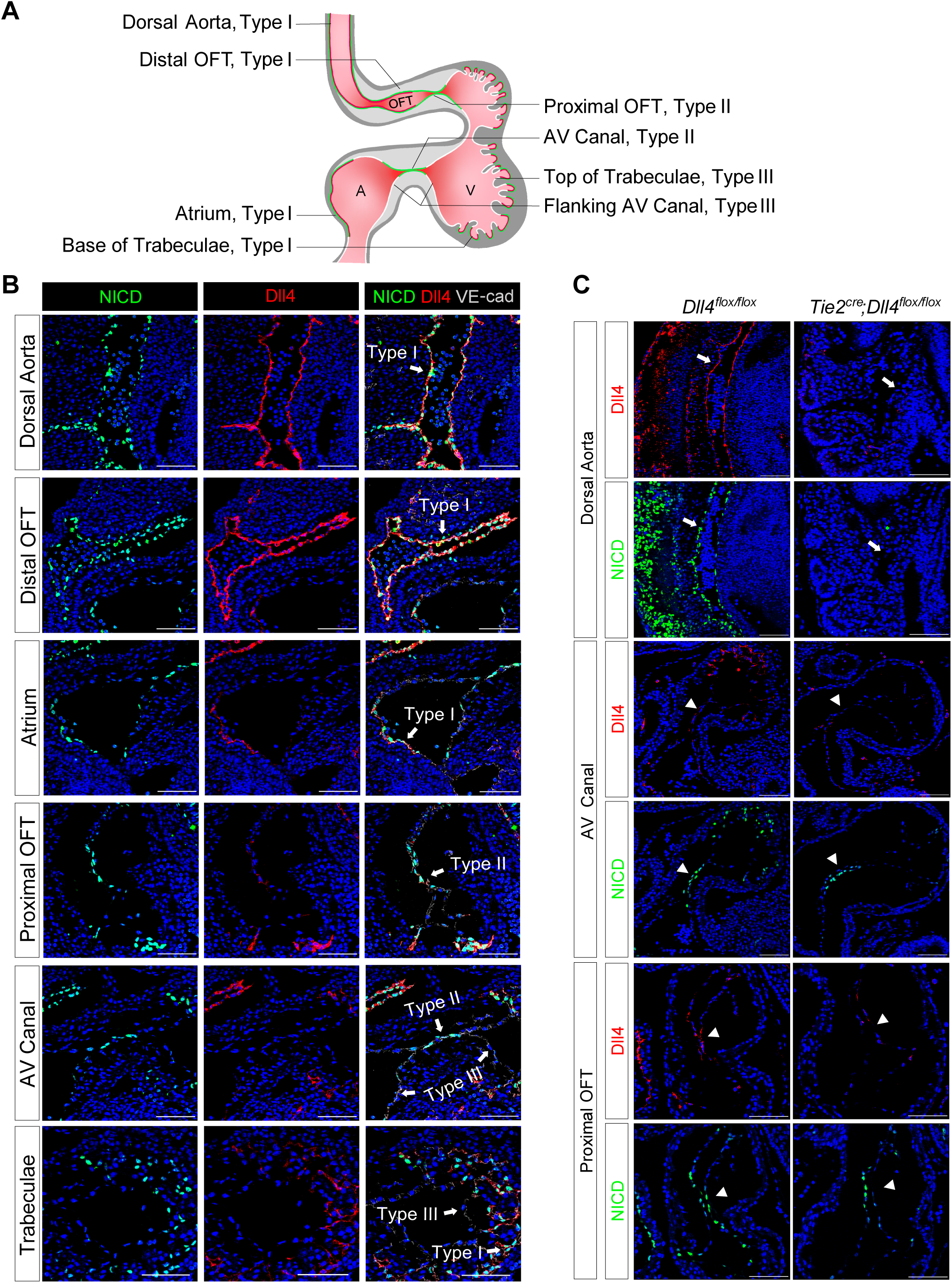
Endocardium and vascular endothelium of dorsal aorta showed different patterns of Notch1 activation and Dll4 ligand expression. **A,** Schematic representation of 3 types of endothelial cells and their corresponding localizations. **B,** Type I cells are NICD-high and Dll4-high. Type II cells are NICD-high and Dll4-low. Type III cells are NICD-low and Dll4-low. Scale bars, 100 µm. (*n* = 3 embryos) **C,** Representative images showing Dll4 protein and NICD in wild-type and endothelial-specific *Dll4*-deleted (*Tie2^cre^;Dll4^flox/flox^*) E9.5 mouse hearts. Scale bars, 100 µm. (wild-type: *n* = 3 embryos; *Tie2^cre^;Dll4^flox/flox^*: *n* = 3 embryos)

One previous study showed that endocardial EMT was dependent on Dll4 ^13^. To further dissect the role of Dll4 in regional Notch activation, we conditionally deleted Dll4 in all endothelial cells using the *Tie2^cre^* line. At E9.5, *Dll4* conditional null AVC cushions were poorly cellularized, consistent with the previous finding. Furthermore, in most areas normally expressing high Dll4, i.e., vascular endothelium and aortic sac, NICD was completely abolished, whereas in the AVC, proximal OFT, and the base of the trabeculae, NICD was reduced but not completely absent (Figure 1C). As conditional deletion of Dll4 resulted in embryos with severely disorganized vasculature (Figure 1C), we cannot rule out the possibility that the reduced NICD in the AVC might not be simply due to Dll4 deletion, but rather affected by circulatory failure. These findings indicate that vascular endothelium is dependent on Dll4 for Notch activation, whereas in the endocardium, additional factors are required to pattern Notch activation in the proximal OFT and AVC.

### 3.2. Notch activation in cushion endocardium is dependent on blood flow

Work in the past on zebrafish embryos showed that cardiac contraction activates endocardial Notch signaling^21^. To test if Notch activation is regulated by blood flow in the developing mouse heart, pregnant mice were gavaged with a single dose of dofetilide at 2mg/kg at E9.5. The embryos were then either harvested immediately after the treatment or allowed to develop until E18.5 when cardiac morphology was assessed (Figure 2—Figure supplement 1A). Dofetilide caused an immediate stop of blood flow in the embryos, as observed by doppler ultrasound from 1 h through to 3 h, which completely recovered at 5 h post-treatment (Figure 2A and Video S5, S6). Three hours cessation of blood flow caused a complete loss of NICD without affecting total Notch1 receptor protein in the proximal OFT and AVC endocardium. The loss of NICD was completely recovered at 5 h post-treatment, in line with the dynamics of the heart rate changes (Figure 2B), while no significant changes in the expressions of Dll4 and Jag1 were noted in AVC endocardium after dofetilide treatment (Figure 2B). Pro-EMT markers phospho-Smad1/5, Sox9 and Twist1 were downregulated in the AVC endocardium after cardiac arrest (Figure 2B). Interestingly, NICD in the dorsal aorta was resistant to the cessation of flow (Figure 2—Figure supplement 1B). Consistent with the inhibition of EMT, transient cessation of blood flow resulted in hypoplastic AVC endocardial cushions 5 hours after treatment (Figure 2—Figure supplement 1C) and more pronounced cushion hypoplasia one day after treatment (Figure 2C), and various heart defects (Figure 2D and Figure 2—Figure supplement 1D) in 40% of embryos: ventricular septal defects (VSD), bicuspid semilunar valve, atrioventricular valve defects, and conotruncal defects, consistent with malformation of endocardial cushions.

**Figure 2:**
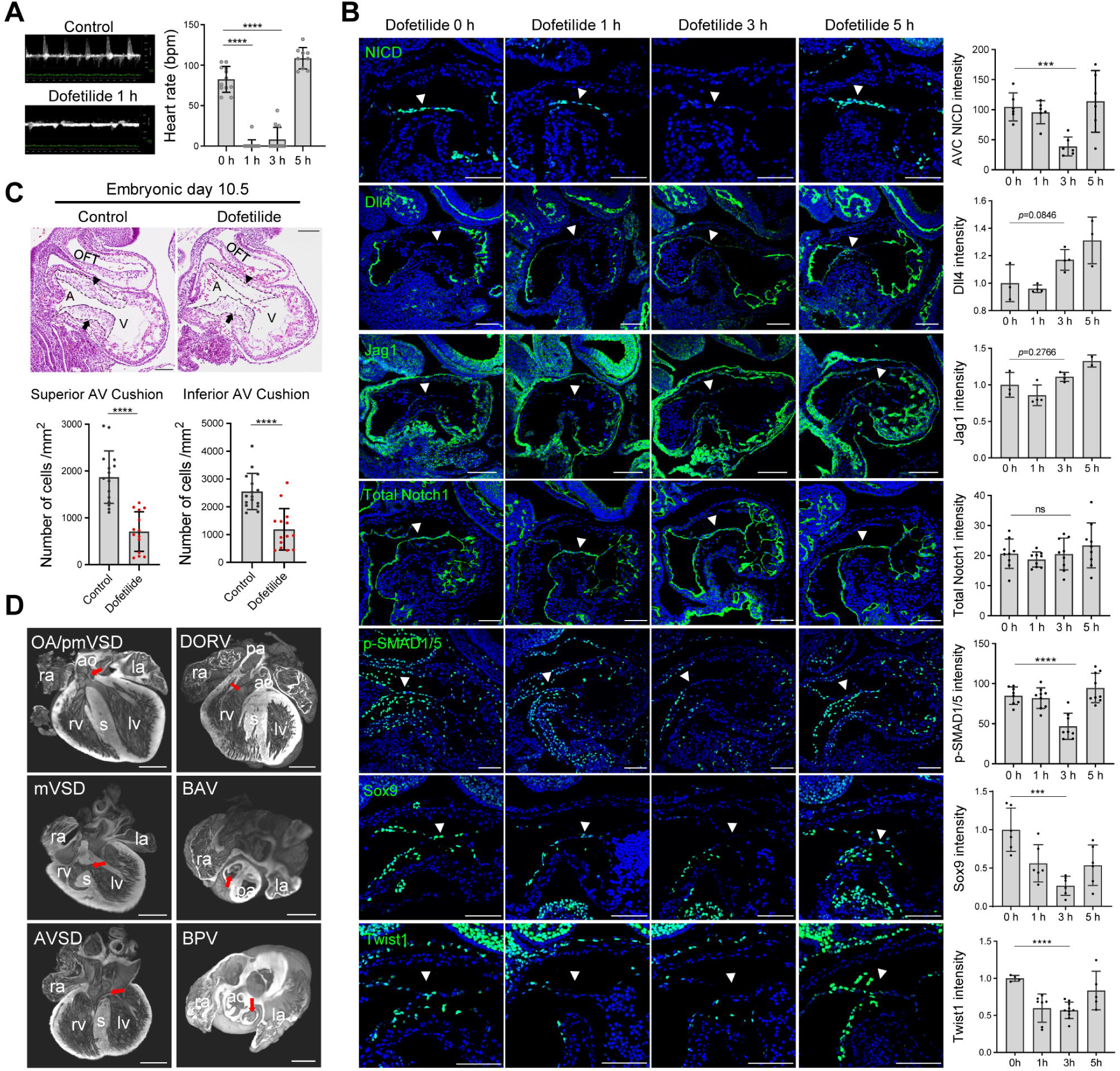
Notch activation in cushion endocardium is dependent on blood flow. **A,** Echocardiography of control and dofetilide treated embryos. (0 h: *n* = 12; 1 h: *n* = 15; 3 h: *n* = 12; 5 h: *n* = 9). **B,** Expression of NICD, Dll4, Jag1, total Notch1, p-SMAD1/5 and Sox9, Twist1 in the E9.5 dorsal aorta endothelium (arrow) and endocardium (arrowhead). **C,** Sagittal E10.5 hematoxylin-eosin stained sections demonstrated hypocellularity in both superior (arrowhead) and inferior AV cushion (arrow) caused by dofetilide treatment. Quantification of mesenchymal cell density in superior (below left) and inferior (below right) AV cushion (Control: *n* = 16 embryos; Dofetilide: *n* = 14 embryos). OFT, outflow tract; A, atrium; V, ventricle. **D,** Representative heart defects induced by maternal dofetilide treatment. pmVSD, perimembranous ventricular septal defect; DORV, double-outlet right ventricle; mVSD, muscular VSD; OA, overriding aorta; BAV, bicuspid aortic valve; AVSD, atrioventricular septal defect; BPV, bicuspid pulmonary valve; ra, right atrium; la, left atrium; ao, aorta; rv, right ventricle; lv, left ventricle; s, interventricular septum; pa, pulmonary artery. Scale bars, 100 µm (**B, C**), 500 µm (**D**). (*n* > 3 for every group and each condition)

To rule out the possible direct effect of dofetilide on endocardial cells independent of flow, we conditionally deleted *Tnnt2* in the embryonic myocardium, which caused bradycardia in the E9.5 embryos and lethality at E10.5 (Video S7). Similar to the effect of dofetilide, reducing blood flow rate by genetic means also caused a significant reduction of NICD in the cushion endocardium at E9.5 (Figure 2—Figure supplement 2A). Furthermore, block of heart beat by *ex vivo* blebbistatin treatment, an inhibitor for non-muscle myosin II ATPase, also prevented Notch cleavage in the cushion endocardium at E9.5 (Figure 2—Figure supplement 2B). To rule out the possible effect of hypoxia on Notch activation, we depleted embryonic erythrocytes by crossing Epor^cre^ with ROSA-DTA mouse line, which resulted in widespread hypoxia without interfering embryonic heartbeat, yet NICD was normal in the AVC endocardium (Figure 2—Figure supplement 2C). We also cultured E9.5 embryos in a medium saturated with 95% O_2_ in the presence of dofetilide for 3 h. High levels of O_2_ eliminated HIF-1α nuclear staining induced by cardiac arrest, while NICD was still absent in the AVC endocardium (Figure 2—Figure supplement 2D). Thus, all subsequent *ex vivo* embryo culture experiments were performed in 95% O_2_ to minimize the effect of hypoxia on Notch activation. Collectively, these results indicate that Notch activation and EMT in the cushion endocardium are dependent on the mechanical stimulation from blood flow.

### 3.3. Flow-responsive mTORC2-PKCε activity is required for Notch activation in the cushion endocardium

Given the fast response of NICD to flow cessation, we suspect a change in post-translational modification, such as a phosphorylation event, that might mediate the effect. PKC, AKT, and ERK have all been reported to have their phosphorylation status altered in response to fluid shear stress in cultured endothelial cells ^22^. In addition, *in vitro* shear stress-induced Notch activation can be blocked by inhibitors of PKC, AKT, and ERK ^23^. Therefore, we examined the phosphorylation status of these three signaling molecules in the endocardium before and after dofetilide treatment. Both pPKC (βII^Ser660^) and pAKT^Ser473^ were restricted to the cushion endocardium and the base of the trabeculae in control embryos and were almost completely lost as quickly as 1 h after treatment and then both recovered at 5 h after treatment. Both pPKC and pAKT were not detectable in dorsal aorta endothelium (Figure 3A and Figure 3—Figure supplement 1A). Their response rate in cushion endocardium was faster than NICD, whose maximum inhibition occurred at 3 h post-treatment (Figure 2C). pERK was not detectable in the cushion endocardium (Figure 3—Figure supplement 1B). Inhibition of AKT phosphorylation in cultured E9.5 embryos by wortmannin did not inhibit Notch activation (Figure 3—Figure supplement 1C), whereas inhibition of PKC activity by staurosporine treatment blocked Notch activation (Figure 3B).

**Figure 3:**
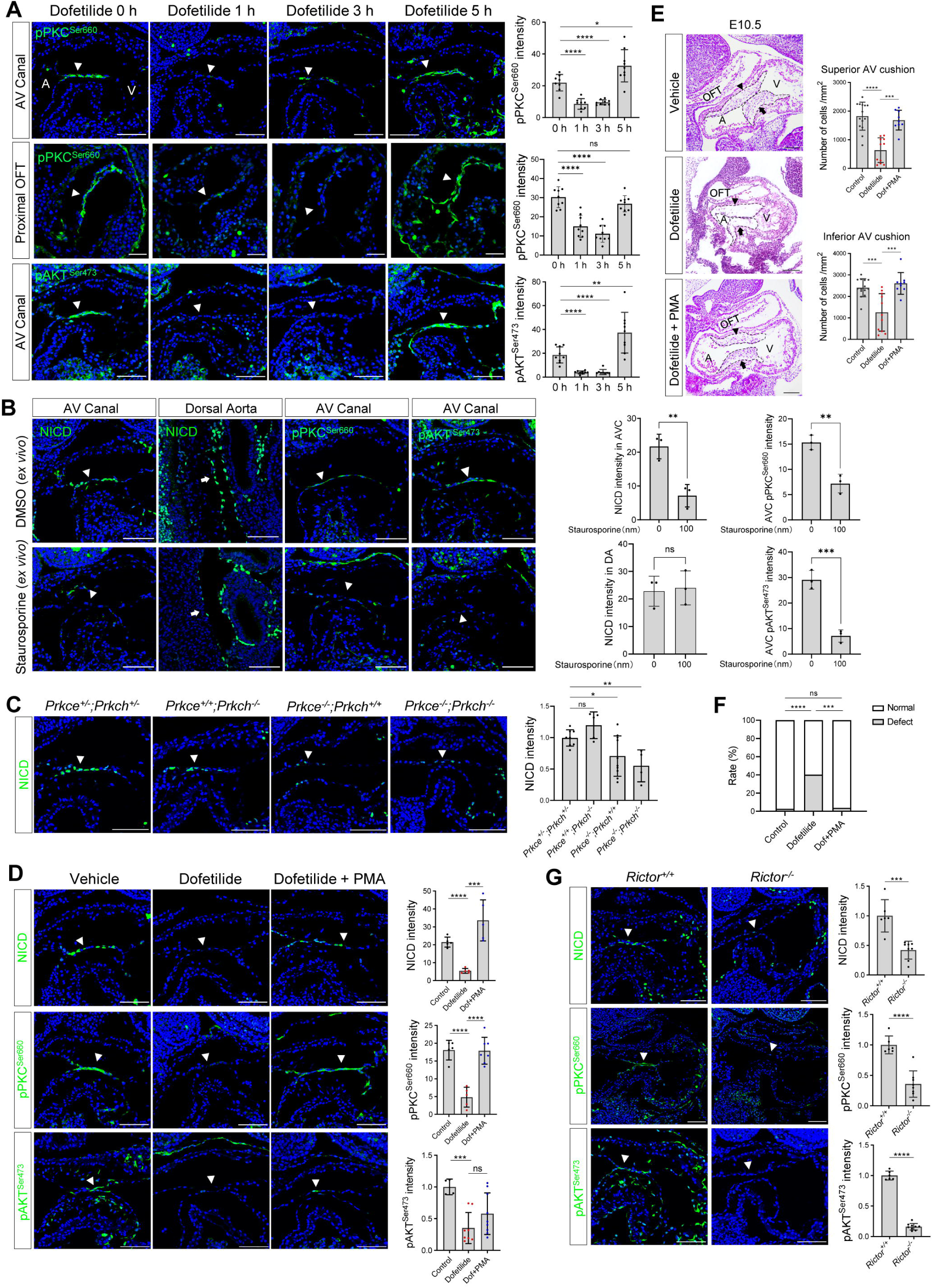
Flow-responsive mTORC2-PKCε activity is required for Notch activation in the cushion endocardium. **A,** Phospho-PKC^Ser660^ and phospho-AKT^Ser473^ levels in the atrioventricular (AV) canal endocardium and proximal OFT endocardium (arrowhead) after dofetilide treatment. Each point in the quantification chart represents one embryo. **B,** NICD, pPKC and pAKT expression in cultured E9.5 heart in response to *ex vivo* staurosporine treatment (100 nM). Each point in the quantification chart represents one embryo. **C,** NICD expression in AV canal endocardium in wild-type, *Prkce* and *Prkch* single knockout and double knockout embryos. Each point in the quantification chart represents one embryo. **D,** NICD, p-PKC^Ser660^ and p-AKT^Ser473^ staining in AVC endocardium (arrowhead) after dofetilide treatment and after rescue by PMA. **E,** Sagittal E10.5 HE staining sections demonstrates acellularized superior (arrowhead) and inferior AV cushion (arrow) caused by dofetilide which were rescued by in PMA treatment (2 mg/kg). OFT, outflow tract; A, atrium; V, ventricle. Mesenchymal cell density was quantitated in superior and inferior AV cushion (Control: *n* = 14 embryos; Dofetilide: *n* = 11 embryos; Dofetilide + PMA: *n* = 9 embryos). **F,** Heart defect rate caused by maternal dofetilide and PMA treatment (Control: *n* = 37; Dofetilide: *n* = 62; Dofetilide + PMA: *n* = 27). **G,** NICD, phospho-PKC^Ser660^, and phospho-AKT^Ser473^ in AV canal endocardium (arrowhead) in wild-type and *Rictor* null mice (*Rictor^+/+^*: *n* = 6; *Rictor^−/−^*: *n* = 8). Scale bars, 100 µm.

To further confirm the role of PKC in regulating Notch, we individually knocked out *Prkce* and *Prkch* (Figure 3—Figure supplement 2C). Among all PKC family members, these two isozymes are most highly expressed in the endocardium based on RNAseq analysis of E9.5 embryos’ endocardial and vascular endothelial cells (Figure 3—Figure supplement 2A). *Prkch*^KO^ did not affect Notch activation or produce any phenotype. *Prkce*^KO^ caused a slight but significant reduction of NICD in the cushion endocardium at E9.5 and caused 25% heart defects at E18.5. *Prkce/Prkch* double knockout resulted in a further loss of NICD at E9.5, leading to pericardial effusion and complete lethality at E10.5. (Figure 3C and Figure 3— Figure supplement 2E). No pericardial effusion or heart beating abnormalities was noted at E9.5 in these mutant embryos (Figure 3—Figure supplement 2D), suggesting a direct action of PKCs on Notch signaling, rather than indirect action through affecting blood flow. Treatment of pregnant mice at E9.5 with a potent PKC activator, phorbol 12-myristate 13-acetate (PMA), almost completely rescued dofetilide-induced loss of NICD, pPKC, cushion hypoplasia, and cardiac defects (Figure 3D-3F). The restricted activation of NICD in AVC region by PMA treatment is consistent with the restricted expression of PKCε and PKCη in the AVC endocardium (Figure 3-supplement figure 2B). Thus, PKC activity is both necessary and sufficient for Notch activation in the cushion endocardium.

Both PKC and AKT belong to the AGC kinase family and both have a conserved hydrophobic motif at the C-terminal tail of the catalytic domain, whose phosphorylation is dependent on mTORC2 and is required for kinase activity ^24^. Therefore, the observed inhibition of hydrophobic motif phosphorylation of PKC and AKT indicates impairment of the common upstream activator mTORC2. We conditionally deleted one of the essential mTORC2 components, Rictor^25^, in the endothelial cells (Figure 3—Figure supplement 2C). As expected, ablation of mTORC2 completely blocked hydrophobic motif phosphorylation of both PKC and AKT and consequently led to significant decrease of NICD in the cushion endocardium (Figure 3G). Thus, blood flow activates Notch in the cushion endocardium via mTORC2-PKC signaling pathway.

### 3.4. Shear stress-induced alteration of membrane lipid microstructure activates mTORC2-PKC-Notch signaling pathway

In cultured mammalian endothelial cells, fluid shear stress (FSS) increases the number of cell surface caveolae, a specialized membrane microdomain enriched for cholesterol and sphingolipids; sequestration of cholesterol inhibits shear-dependent activation of ERK ^26^. Thus, we tried to manipulate the membrane lipid microdomain by treating the cultured E9.5 embryos with cholesterol. Staining of Caveolin-1, the major component of caveolae, showed that Caveolin-1 was normally expressed on the cell surface of the AVC endocardial cells including the luminal surface and the lateral cell adhesion sites. However, in area with low shear stress such as dorsal aorta endothelium, atrial, and ventricular endocardium downstream of AVC, Caveolin-1 was mainly restricted to the lateral cell adhesion sites and absent on the luminal surface. *Ex vivo* dofetilide treatment caused retraction of Caveolin-1 from the luminal surface to the lateral cell adhesion sites in the AVC endocardial cells, while co-treatment with cholesterol rescued the presence of Caveolin-1 to the luminal surface of AVC endocardial cell in the presence of dofetilide (Figure 4A). In addition, scanning electron microscopy on E9.5 AV canal endocardium showed numerous membrane invaginations on the luminal surface of the endocardial cells. The size of the invaginations ranged from 50 to 100 nm, consistent with the reported size of caveolae. Dofetilide significantly reduced the number of membrane invaginations, which recovered after restore of blood flow at 5 hours post dofetilide treatment (Figure 4—Figure supplement 1A). The reduction of membrane invaginations at 3 hours post ex vivo dofetilide treatment could be rescued by co-treatment of cholesterol (Figure 4B). The loss of NICD, pPKC^Ser660^, and pAKT^Ser473^ caused by dofetilide could all be rescued by pretreatment with cholesterol, indicating reactivation of mTORC2 (Figure 4C). These rescuing effects of cholesterol disappeared in the *Rictor* KO embryos (Figure 4C), suggesting mTORC2 was intermediate signaling molecules linking membrane lipid microstructure and Notch activation.

**Figure 4:**
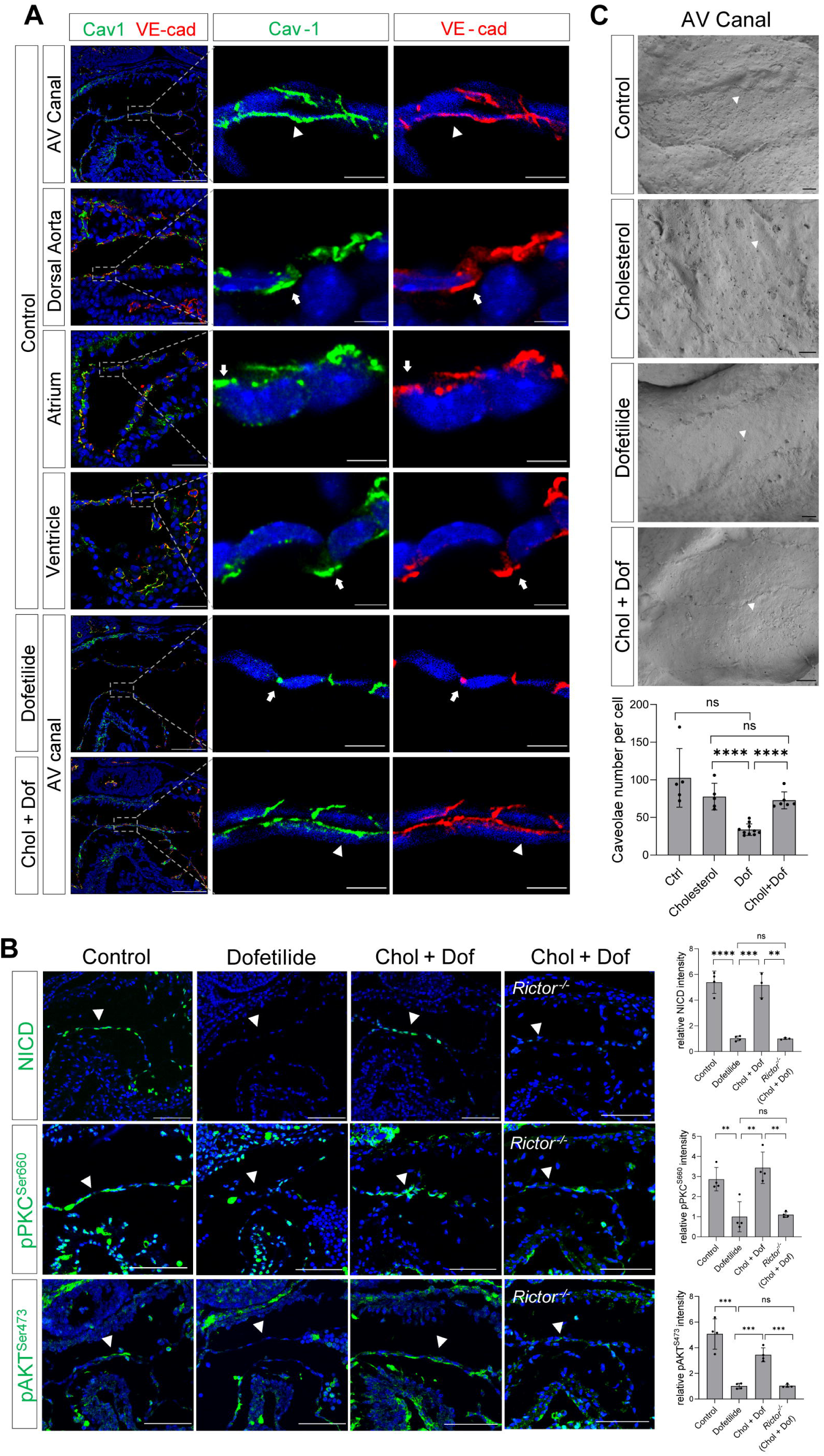
Shear stress-induced alteration of membrane lipid microstructure activated mTORC2-PKC-Notch signaling pathway. **A,** Caveolin-1 and VE-cadherin expression in mouse E9.5 AV canal, dorsal aorta, atrium and ventricle endocardium, demonstrating luminal (arrowhead) and lateral (arrow) surface localization. *Ex vivo* dofetilide treatment (0.2 μg/ml) of cultured E9.5 embryos for 3 h caused retraction of Caveolin-1 and VE-cadherin from luminal surface of the AVC endocardial cells to the lateral cell adhesion sites which could be rescued by co-treatment with cholesterol (1 mg/ml). **B,** Representative images and quantifications of scanning electron microscopy on E 9.5 embryonic heart AV canal endocardial cells at their luminal surfaces, with caveolae structure presentively pointed by arrowhead. Each point in the quantification chart represents one embryo. **C,** Loss of NICD, phospho-PKC^Ser660^, and phospho-AKT^Ser473^ in AVC endocardium (arrowhead) by *ex vivo* dofetilide treatment could be rescued by cholesterol. The rescue failed in *Rictor* null hearts. Each point in the quantification chart represents one embryo.Scale bars, 100 µm (**A, C**). Scale bars, 10 µm (**A**, zoom-in), 1 µm (**B**).

### 3.5. Pharmacogenetic interaction in etiology of congenital heart defects

Gene-environmental interactions are believed to underlie the complex etiology of many congenital heart diseases. With the mechanosensitive nature of Notch in mind, we speculated that reducing the gene dosage of Notch1 may interact with agents that disrupt normal hemodynamic stimulations to the endocardium and synergistically causes heart defects. To test this, we crossed Notch1 heterozygous null male (FVB) with wildtype female mice, and treated the pregnant mice with a lower dose of dofetilide (1.8 mg/kg) at E9.5. In addition, we tested another drug verapamil, a commonly used FDA-approved L-type calcium channel blocker for treatment of high blood pressure, heart arrhythmias, and angina^27^, in FVB background. A single dose of maternal verapamil treatment at E9.5 significantly decreased embryonic heart rate (Figure 5E). Neither Notch1 heterozygosity alone, nor drug treatment alone at the applied dosage produced any notable heart defects. However, the combination of Notch1 mutation and drug exposure of either dofetilide or verapamil resulted over 50% of fetuses having various heart defects of endocardial origin (Figure 5A-D).

**Figure 5:**
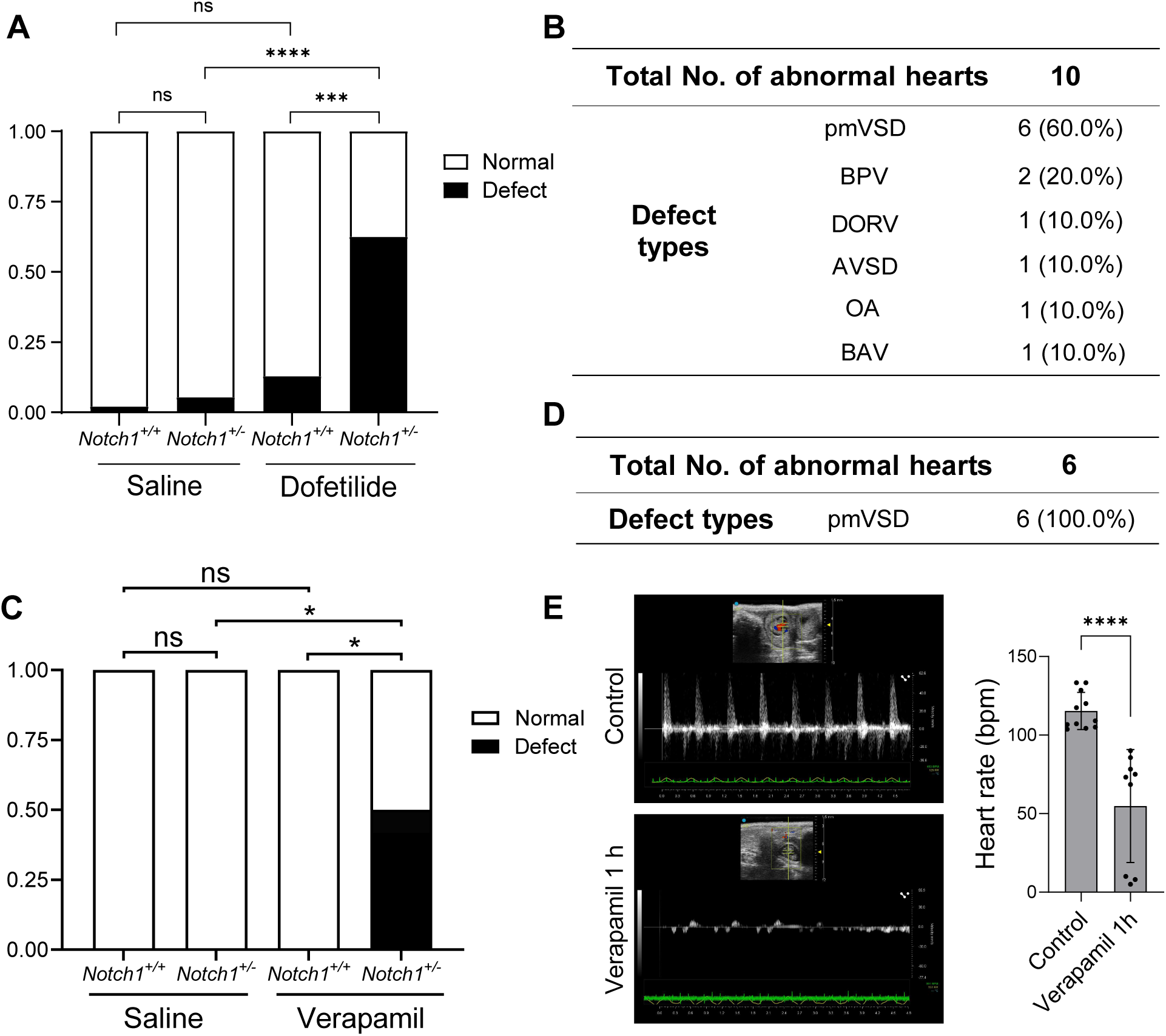
Pharmacogenetic interaction causing heart defects. **A,** Heart defect rate significantly increased in Notch1 heterozygous embryos treated with dofetilide (1.8 mg/kg) at E9.5. (Saline *Notch1^+/+^*: *n* = 48, Saline *Notch1^+/−^*: *n* = 37, Dofetilide *Notch1^+/+^*: *n* = 31, Dofetilide *Notch1^+/−^*: *n* = 16). **B,** Types of heart defects in the dofetilide and *Notch1^+/−^* combined group. **C,** Heart defect rate significantly increased in Notch1 heterozygous embryos treated with Verapamil (15 mg/kg) at E9.5. (Saline *Notch1^+/+^*: *n* = 18, Saline *Notch1^+/−^*: *n* = 9, Verapamil *Notch1^+/+^*: *n* = 8, Verapamil *Notch1^+/−^*: *n* = 12). **D,** Types of heart defects in the verapamil combined with *Notch1^+/−^* group. **E,** Representative echocardiography and quantifications of heartbeat in control and verapamil treated E 9.5 embryos. Each point in the quantification chart represents one embryo.

## 4. DISCUSSION

In this study, we identified fluid shear stress as the primary activator of Notch signaling in the area of AVC and OFT of the mouse looping heart tube. Strong fluid shear stress in the AVC and OFT enhances PKC phosphorylation by mTORC2 possibly by maintaining a particular membrane microstructure. Activated PKC then augments Notch cleavage under the minimal ligand stimulation, resulting Notch activation and EMT specifically in areas of valve primordium characteristic of narrow internal diameter and high shear stress. These findings are directly relevant for endocardial patterning. Notch activity is known to be essential for establishing OFT and AVC endocardial cell identity capable of EMT ^10, 11^. Activation of Notch in the AVC endocardium is presumed to be mainly driven by the Dll4 ligand because Dll4 has been demonstrated to be expressed in the AVC endocardial cells and endothelial-specific knockout of *Dll4* produced acellularized AV cushions ^13^. However, careful examination of the immunostaining pattern revealed that NICD, VEGFR2 and Dll4 were initially expressed with equal levels throughout cardiovascular endothelium at E8.5 prior to endocardial EMT, but at E9.5 with the onset of EMT, endocardial VEGFR2 and Dll4 were markedly reduced and NICD becomes restricted to endocardium overlaying the merging OFT and AVC cushions, implying a transition of the main driver of Notch activation from the pan-endothelial VEGFR2-Dll4 stimulation at E8.5 to the regionally restricted shear force stimulation at E9.5.

We therefore propose the following working model as how mechanical cues guide Notch activation during endocardial patterning. Following gastrulation, high level of VEGF-VEGFR2 signaling in mesodermal tissues initiates Dll4 expression which activates Notch and drives arterial endothelium and endocardium differentiation and proliferation. At E9.5 when endocardial EMT starts, VEGF signaling must dampen due to its suppressive function on EMT ^28, 29^. One way to lower VEGF signaling in the endocardium is through downregulating VEGFR2 expression. Consequently, Dll4 expression also decreases, rendering Notch less active in the endocardium compared to the vascular endothelium. In the meantime, cardiac ballooning on both sides of the AVC expands the myocardial chambers and widen the endocardial internal diameter, leaving only the endocardium of the OFT and AVC endocardial cushion region closely attached. Narrow blood passage way creates strong shear stress to stimulate Notch cleavage only in these narrow areas of future cardiac septation and valve formation. Thus, in the developing mouse hearts: (1) VEGF signaling is reduced to permit endocardial EMT; (2) Dll4 expression is reduced to prevent widespread endocardial Notch activation and make endocardium sensitive to flow; (3) a proper cushion size and shape is maintained by limiting the flanking endocardium to undergo EMT despite physically close to the field of BMP2 derived from of AVC myocardium (Figure 6).

**Figure 6.**
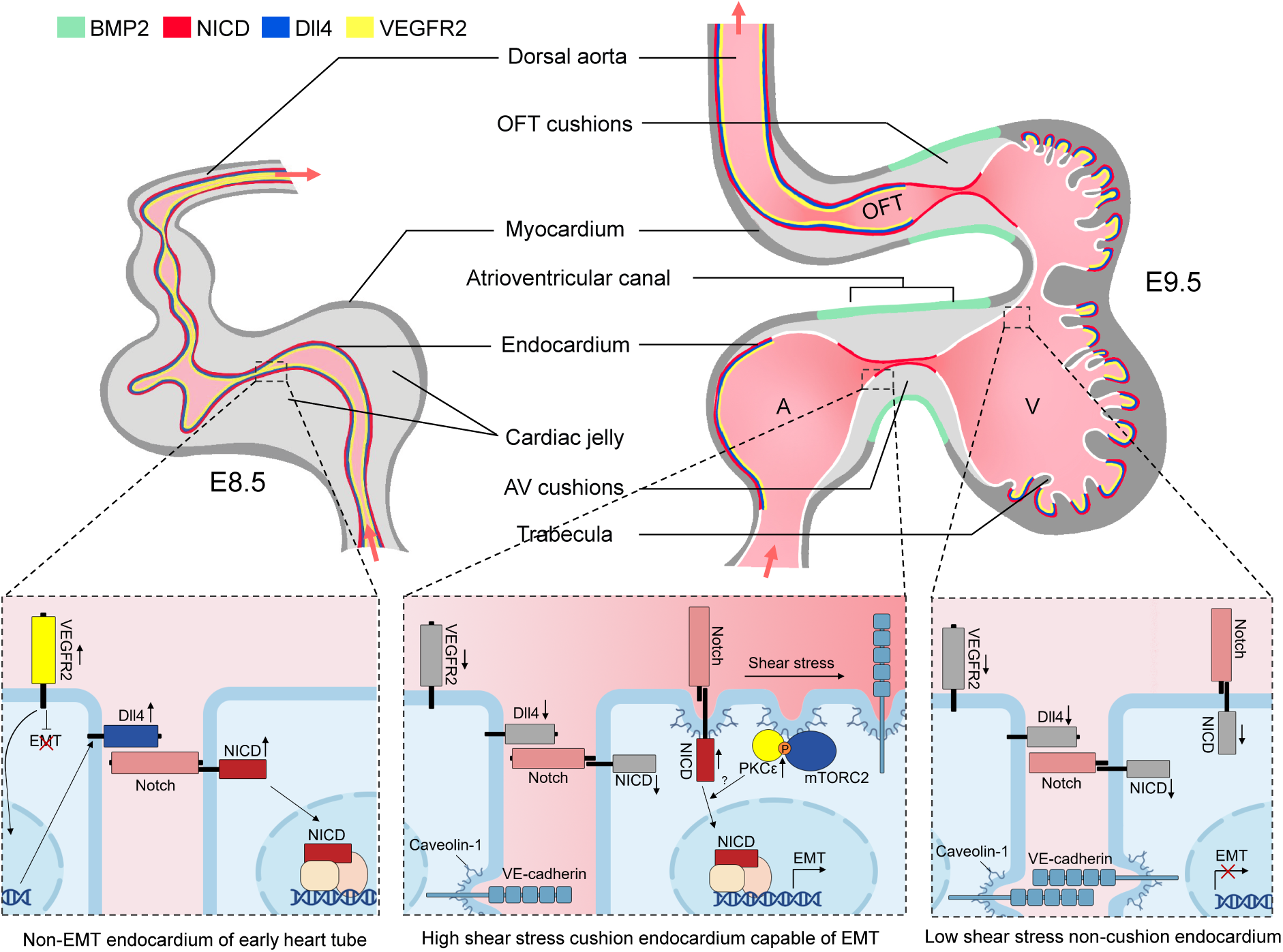
Working model of the establishment of Notch activation pattern by mechanical cues. The establishment of Notch activation pattern by mechanical cues involves a series of events in the developing heart tube. At E8.5, the arterial endothelium and non-EMT endocardium exhibit low shear stress, high VEGF, Dll4, and Notch signaling. One day later, the endocardium undergoes patterning and becomes capable of epithelial-to-mesenchymal transition (EMT) only in the AVC and proximal OFT regions. This patterning is achieved through restricted expression of BMP2 and NICD in these specific areas. EMT requires the downregulation of VEGF signaling by the endocardium, enabling EMT to occur. Additionally, Dll4 is downregulated in endocardium to prevent widespread Notch activation. Simultaneously, the high shear stress present in the AVC and proximal OFT regions leads to increased membrane lipid order, which activates the mTORC2-PKC-Notch pathway and promotes EMT. On the other hand, regions flanking these valve-forming areas experience lower shear stress, resulting in inactive Notch signaling and an inability to undergo EMT.

Studies in zebrafish identified several cation channels including Trpv4, Trpp2, Piezo1, Piezo2 and P2X that contribute to mechanosensing in the developing endocardium ^15, 17, 30^. Blood flow in the looping mammalian heart is predominantly unidirectional whereas early zebrafish embryonic heart displays oscillatory flow pattern ^17^. Mammalian embryonic endocardium undergoes extensive EMT to form valve primordia while zebrafish atrioventricular valve primordia is formed via partial EMT and the collective cell migration of endocardial cells into the cardiac jelly followed by tissue sheet delamination^15, 31, 32^. It is thus possible that mammalian hearts may evolve additional mechanisms to guide endocardial patterning and development. Our data support a model that membrane lipid microdomain acts as a shear stress sensor and transduces the mechanical cue to activate intracellular mTORC2-PKC-Notch signaling pathway in the developing endocardium. Shear stress has been shown to upregulate the number of caveolae in cultured endocardial cells ^26^. Flow induced AKT phosphorylation is dependent on Caveolin-1 ^33^. Both PKCα and PKCε interact with Caveolin-1 and bind with caveolae membranes ^34, 35^. In neuronal cells or neural tissues, accumulation of psychosine in the plasma membrane disrupted lipid rafts, blocked recruitment of mTORC2 and PKC to the lipid raft, and inhibited AKT and PKC activation ^36, 37^. In line with these previous findings, we demonstrated that high level of shear stress in the AVC and OFT is required for caveolin localization to the luminal surface of the endocardial cell membrane, and this membrane lipid microstructure is both necessary and sufficient for mTORC2-PKC-Notch pathway activation. The mechanism by which shear stress and cholesterol increases caveolae is unknown. In a previous study, shear stress exposure for a few minutes rapidly decreases the lipid order and increase the fluidity of the plasma membranes which appears to contradict our findings ^38^. However, membrane lipid compositions are highly dynamic and regulated. It is possible that acute shear stress and its associated kinetic energy may initially decrease membrane lipid order, but long-term shear may evoke cellular feedback mechanisms to increase lipid order and enhance membrane rigidity. As cholesterol is an integral component of lipid raft and caveolae, it is likely that enrichment of cholesterol to the plasma membrane by exogenous supplementation might alter the membrane structure to make the lipid raft structure less dependent on sheer stress. It is noteworthy that the methodology used to alter blood flow in this study inevitably affects myocardial contraction. Thus, further work to uncouple changes in shear stress and myocardial mechanical properties, with the aid of theoretical modeling or using mouse heart valve explants, is needed to fully characterize the effect of shear stress on mouse endocardial development.

The positive regulation of Notch activation by PKC has been reported in a number of *in vitro* systems. Studies in CD4^+^ T cells revealed that Notch is activated within hours of TCR stimulation independent of Notch ligands, and PKC activity is both sufficient and necessary for the ligand independent Notch activation ^39^. The exact mechanism by which PKC regulate Notch processing is not known, and in need of further investigation in the future.

The vast majority of congenital heart diseases do not have a genetic explanation and are considered to have a multifactorial origin. Studies conducted on various model organisms such as chick and zebrafish have demonstrated that alterations in blood flow during early stages of heart development can lead to heart malformation ^40^. Similarly, human studies have shown that fetal bradycardia or abnormal flow patterns in the Ductus Venosus during the first trimester of pregnancy are associated with an increased risk of CHD ^41, 42^. Our research findings suggest that abnormal heart contraction or flow patterns, resulting from genetic mutations or the use of certain drugs during early heart development, may interact with genetic pathways involved in mechanosensing and endocardial EMT. These interactions could contribute to the complex etiology of congenital heart disease.

## 5. ACKNOWLEDGEMENTS

The authors are grateful to Dr. Xu Li of Westlake University for valuable suggestions regarding Notch experiments. We thank Dr. Zhen Zhang from Shanghai Jiao Tong University, Dr. Bing Zhang, Dr. Jiemin Jia, Dr. Shang Cai from Westlake University and Dr Hui Zhang from Shanghai Tech University for sharing mouse lines. We thank the Microscopy Core Facility of Westlake University for assistance and advice. We thank the Westlake Animal Facility and Youshi Chen for mouse breeding and husbandry. We thank Jianfeng Wang for assistance in *ex vivo* culture. This work was supported by Natural Science Foundation of Zhejiang Province of China (LZ19H040001) and Westlake Education Foundation.

## 6. AUTHOR CONTRIBUTIONS

H.S. raised hypothesis, designed experiments, interpreted data and drafted the work. Y.M., and S.H. conducted the main body of the experiments, including mouse crosses and treatment, embryo harvest, staining and data analysis. X.L. made contributions by optimizing light sheet microscopy and performed work related to heart phenotyping. S.H. participated in manuscript drafting. X.T. characterized the change of membrane lipid structure induced by flow. J.L. assisted in the performances of echocardiography for both pregnant and fetal mice.

## Extended Data

**Figure 1—Figure supplement 1: Notch is strongly activated in the AVC and proximal OFT endocardium despite weak ligand expression at the onset of EMT. A,** Representative images of NICD (green) whole mount immunofluorescence, *Dll4* (red) whole mount *in situ* hybridization, and Jag1 (yellow) immunofluorescence of E8.5 and E9.5 mouse heart. Arrows indicate arterial endothelium and arrowheads indicate endocardium. **B,** Representative images demonstrating VEGFR2 (Green) and Dll4 (red) protein expression in E8.5 and E9.5 mouse artery (arrowhead) and endocardium (arrow). **C,** Quantification of NICD and Dll4 immunofluorescence intensity within the area of each 3 types of endothelial cells.

**Figure 2—Figure supplement 1. A,** Schematic diagram of the experimental design. **B,** Representative figures and statistics of NICD expression in dorsal aorta and POFT after dofetilide treatment. **C,** HE staining of E9.5 embryonic hearts and the quantifications of superior and inferior AV cushion cell density in control group and 5 hours after dofetilide treatment. (Control: *n* = 7 embryos; Dofetilide: *n* = 6 embryos) **D,** Heart defects induced by maternal dofetilide treatment. (0 mg/kg: n = 37; 2.0 mg/kg: n =117). pmVSD, perimembranous ventricular septal defect; DORV, double-outlet right ventricle; mVSD, muscular VSD; OA, overriding aorta; BAV, bicuspid aortic valve; AVSD, atrioventricular septal defect; BPV, bicuspid pulmonary valve.

**Figure 2—Figure supplement 2. A,** Representative images of NICD in myocardial-specific cardiac troponin T knockout mouse heart. **B,** Representative images of NICD expression in AV canal, proximal OFT and dorsal aorta after blebbistatin treatment (5 µM) for 3 h. **C,** HIF-1α and NICD in red blood cell depleted E9.5 mouse hearts. **D,** Representative images of HIF-1α and NICD in E9.5 control, dofetilide-treated, and hyperoxia-treated mouse hearts. Arrow: dorsal aorta endothelium. Arrowhead: endocardium. Scale bars, 100 µm.

**Figure 3—Figure supplement 1. A,** pPKC^Ser660^ in E9.5 mouse dorsal aorta, and pAKT^Ser473^ in E9.5 dorsal aorta endothelium. **B,** pERK1/2 was not detectable in AV canal endocardium. **C,** NICD in cultured E9.5 heart in response to *ex vivo* wortmannin treatment (2 μM).

**Extended Figure 3—Figure supplement 2. A,** Counts of conventional and novel PKC isoforms assessed by bulk RNA-seq of E9.5 embryonic endocardial cells. (n =4) **B,** Immunofluorescence staining of PKCε and PKCη in E9.5 embryonic hearts. **C,** *Prkch*, *Prkce* and *Rictor* knockout mice strategies and western blot or immunofluorescence confirmation of KO. **D,** Echocardiography of heterozygous *Prkch*, *Prkce* knockouts. **E,** Counts of abnormal hearts of the descendants of *Prkch*, *Prkce* double heterozygous intercrosses at E18.5. Arrow: dorsal aorta endothelium. Arrowhead: endocardium. Scale bars, 100 µm.

**Extended Figure 4—Figure supplement 1. A,** Scanning electron microscopy of E9.5 embryonic heart in AV canal endocardial cells at their luminal surfaces after dofetilide treatment at 3 hours and 5 hours, with caveolae structure presentively pointed by arrowhead. Caveolae density was quantified. Each group has 4 embryos.

**Table S1: Materials and Reagents.**

**Video S1: E8.5 NICD whole mount staining.** NICD (green) whole-mount immunofluorescence staining of E8.5 mouse heart.

**Video S2: E9.5 NICD whole mount staining.** NICD (green) whole-mount immunofluorescence staining of E9.5 mouse heart.

**Video S3: E8.5 Dll4 *in situ* hybridization.** Whole-mount *Dll4* (red) *in situ* hybridization staining of E8.5 mouse heart.

**Video S4: E9.5 Dll4 *in situ* hybridization.** Whole-mount *Dll4* (red) *in situ* hybridization staining of E9.5 mouse heart.

**Video S5: E9.5 echo of a beating embryonic heart.** Echocardiography of a normal E9.5 mouse embryo *in utero* using color Doppler.

**Video S6: E9.5 echo of a dofetilide treated embryonic heart.** Echocardiography of a 1-hour after dofetilide treated E9.5 mouse embryo *in utero* using color Doppler.

**Video S7: E9.5 echo of *cTnT^cre^* x *Tnnt2 ^flox/flox^* beating with non-beating.** Echocardiography of two E9.5 embryos from *cTnT^cre^;Tnnt2 ^flox/+^* x *Tnnt2 ^flox/flox^* crossing. One embryo was genotyped to be *Tnnt2^flox/flox^* and the other *cTnT^cre^; Tnnt2^flox/flox^*.

## REFERENCES

1. Moorman AF and Christoffels VM. Cardiac chamber formation: development, genes, and evolution. Physiological reviews. 2003;83:1223–67.

2. Miquerol L and Kelly RG. Organogenesis of the vertebrate heart. Wiley Interdisciplinary Reviews: Developmental Biology. 2013;2:17–29.

3. Leckband DE, Le Duc Q, Wang N and De Rooij J. Mechanotransduction at cadherin-mediated adhesions. Current Opinion in Cell Biology. 2011;23:523–530.

4. Ma L, Lu M-F, Schwartz RJ and Martin JF. Bmp2 is essential for cardiac cushion epithelial-mesenchymal transition and myocardial patterning. Development. 2005;132:5601–5611.

5. Rivera-Feliciano J and Tabin CJ. Bmp2 instructs cardiac progenitors to form the heart-valve-inducing field. Developmental biology. 2006;295:580–588.

6. Shirai M, Imanaka-Yoshida K, Schneider MD, Schwartz RJ and Morisaki T. T-box 2, a mediator of Bmp-Smad signaling, induced hyaluronan synthase 2 and Tgfbeta2 expression and endocardial cushion formation. Proceedings of the National Academy of Sciences of the United States of America. 2009;106:18604–9.

7. Niessen K, Fu Y, Chang L, Hoodless PA, McFadden D and Karsan A. Slug is a direct Notch target required for initiation of cardiac cushion cellularization. Journal of Cell Biology. 2008;182:315–325.

8. Del Monte G, Grego-Bessa J, Gonzalez-Rajal A, Bolos V and De La Pompa JL. Monitoring Notch1 activity in development: evidence for a feedback regulatory loop. Developmental dynamics : an official publication of the American Association of Anatomists. 2007;236:2594–614.

9. Grego-Bessa J, Luna-Zurita L, Del Monte G, Bolós V, Melgar P, Arandilla A, Garratt AN, Zang H, Mukouyama Y-S, Chen H, Shou W, Ballestar E, Esteller M, Rojas A, Pérez-Pomares JM and De La Pompa JL. Notch Signaling Is Essential for Ventricular Chamber Development. Developmental cell. 2007;12:415–429.

10. Timmerman LA, Grego-Bessa J, Raya A, Bertran E, Perez-Pomares JM, Diez J, Aranda S, Palomo S, McCormick F, Izpisua-Belmonte JC and de la Pompa JL. Notch promotes epithelial-mesenchymal transition during cardiac development and oncogenic transformation. Genes & development. 2004;18:99–115.

11. Luna-Zurita L, Prados B, Grego-Bessa J, Luxan G, del Monte G, Benguria A, Adams RH, Perez-Pomares JM and de la Pompa JL. Integration of a Notch-dependent mesenchymal gene program and Bmp2-driven cell invasiveness regulates murine cardiac valve formation. The Journal of clinical investigation. 2010;120:3493–507.

12. Krebs LT, Xue Y, Norton CR, Shutter JR, Maguire M, Sundberg JP, Gallahan D, Closson V, Kitajewski J, Callahan R, Smith GH, Stark KL and Gridley T. Notch signaling is essential for vascular morphogenesis in mice. Genes & development. 2000;14:1343–52.

13. MacGrogan D, D’Amato G, Travisano S, Martinez-Poveda B, Luxan G, Del Monte-Nieto G, Papoutsi T, Sbroggio M, Bou V, Gomez-Del Arco P, Gomez MJ, Zhou B, Redondo JM, Jimenez-Borreguero LJ and de la Pompa JL. Sequential Ligand-Dependent Notch Signaling Activation Regulates Valve Primordium Formation and Morphogenesis. Circulation research. 2016;118:1480–97.

14. Luxan G, D’Amato G, MacGrogan D and de la Pompa JL. Endocardial Notch Signaling in Cardiac Development and Disease. Circulation research. 2016;118:e1–e18.

15. Duchemin AL, Vignes H and Vermot J. Mechanically activated piezo channels modulate outflow tract valve development through the Yap1 and Klf2-Notch signaling axis. eLife. 2019;8.

16. Galvez-Santisteban M, Chen D, Zhang R, Serrano R, Nguyen C, Zhao L, Nerb L, Masutani EM, Vermot J, Burns CG, Burns CE, Del Alamo JC and Chi NC. Hemodynamic-mediated endocardial signaling controls in vivo myocardial reprogramming. eLife. 2019;8.

17. Heckel E, Boselli F, Roth S, Krudewig A, Belting HG, Charvin G and Vermot J. Oscillatory Flow Modulates Mechanosensitive klf2a Expression through trpv4 and trpp2 during Heart Valve Development. Current biology : CB. 2015;25:1354–61.

18. Mounsey JP and Dimarco JP. Dofetilide. Circulation. 2000;102:2665–2670.

19. D’Amato G, Luxan G, del Monte-Nieto G, Martinez-Poveda B, Torroja C, Walter W, Bochter MS, Benedito R, Cole S, Martinez F, Hadjantonakis AK, Uemura A, Jimenez-Borreguero LJ and de la Pompa JL. Sequential Notch activation regulates ventricular chamber development. Nature cell biology. 2016;18:7–20.

20. Joshua, Lan W, Boudreau E, Stanley, He D, Schachterle W, Didier, Oettgen P, Brian, Benoit and Jason. ETS Factors Regulate Vegf-Dependent Arterial Specification. Developmental cell. 2013;26:45–58.

21. Samsa LA, Givens C, Tzima E, Stainier DY, Qian L and Liu J. Cardiac contraction activates endocardial Notch signaling to modulate chamber maturation in zebrafish. Development. 2015;142:4080–91.

22. Li YS, Haga JH and Chien S. Molecular basis of the effects of shear stress on vascular endothelial cells. Journal of biomechanics. 2005;38:1949–71.

23. Masumura T, Yamamoto K, Shimizu N, Obi S and Ando J. Shear stress increases expression of the arterial endothelial marker ephrinB2 in murine ES cells via the VEGF-Notch signaling pathways. Arteriosclerosis, thrombosis, and vascular biology. 2009;29:2125–31.

24. Baffi TR, Lorden G, Wozniak JM, Feichtner A, Yeung W, Kornev AP, King CC, Del Rio JC, Limaye AJ, Bogomolovas J, Gould CM, Chen J, Kennedy EJ, Kannan N, Gonzalez DJ, Stefan E, Taylor SS and Newton AC. mTORC2 controls the activity of PKC and Akt by phosphorylating a conserved TOR interaction motif. Sci Signal. 2021;14.

25. Sarbassov DD, Ali SM, Kim DH, Guertin DA, Latek RR, Erdjument-Bromage H, Tempst P and Sabatini DM. Rictor, a novel binding partner of mTOR, defines a rapamycin-insensitive and raptor-independent pathway that regulates the cytoskeleton. Current biology : CB. 2004;14:1296–302.

26. Park H, Go Y-M, John PLS, Maland MC, Lisanti MP, Abrahamson DR and Jo H. Plasma Membrane Cholesterol Is a Key Molecule in Shear Stress-dependent Activation of Extracellular Signal-regulated Kinase. Journal of Biological Chemistry. 1998;273:32304–32311.

27. Fahie S and Cassagnol M. Verapamil *StatPearls* Treasure Island (FL) ineligible companies. Disclosure: Manouchkathe Cassagnol declares no relevant financial relationships with ineligible companies.: StatPearls Publishing Copyright © 2023, StatPearls Publishing LLC.; 2023.

28. Dor Y, Camenisch TD, Itin A, Fishman GI, McDonald JA, Carmeliet P and Keshet E. A novel role for VEGF in endocardial cushion formation and its potential contribution to congenital heart defects. Development. 2001;128:1531–1538.

29. Chang C-P, Neilson JR, Bayle JH, Gestwicki JE, Kuo A, Stankunas K, Graef IA and Crabtree GR. A Field of Myocardial-Endocardial NFAT Signaling Underlies Heart Valve Morphogenesis. Cell. 2004;118:649–663.

30. Fukui H, Chow RW, Xie J, Foo YY, Yap CH, Minc N, Mochizuki N and Vermot J. Bioelectric signaling and the control of cardiac cell identity in response to mechanical forces. Science. 2021;374:351–354.

31. Paolini A, Fontana F, Pham V-C, Rödel CJ and Abdelilah-Seyfried S. Mechanosensitive Notch-Dll4 and Klf2-Wnt9 signaling pathways intersect in guiding valvulogenesis in zebrafish. Cell reports. 2021;37:109782.

32. Chow RW-Y, Fukui H, Chan WX, Tan KSJ, Roth S, Duchemin A-L, Messaddeq N, Nakajima H, Liu F, Faggianelli-Conrozier N, Klymchenko AS, Choon Hwai Y, Mochizuki N and Vermot J. Cardiac forces regulate zebrafish heart valve delamination by modulating Nfat signaling. PLoS biology. 2022;20:e3001505.

33. Albinsson S, Nordström I, Swärd K and Hellstrand P. Differential dependence of stretch and shear stress signaling on caveolin-1 in the vascular wall. American Journal of Physiology-Cell Physiology. 2008;294:C271–C279.

34. Mineo C, Ying Y-S, Chapline C, Jaken S and Anderson RGW. Targeting of Protein Kinase Cα to Caveolae. Journal of Cell Biology. 1998;141:601–610.

35. Oka N, Yamamoto M, Schwencke C, Kawabe J-I, Ebina T, Ohno S, Couet J, Lisanti MP and Ishikawa Y. Caveolin Interaction with Protein Kinase C. Journal of Biological Chemistry. 1997;272:33416–33421.

36. Sural-Fehr T, Singh H, Cantuti-Catelvetri L, Zhu H, Marshall MS, Rebiai R, Jastrzebski MJ, Givogri MI, Rasenick MM and Bongarzone ER. Inhibition of IGF-1-PI3K-Akt-mTORC2 in lipid rafts increases neuronal vulnerability in a genetic lysosomal glycosphingolipidosis. Disease models & mechanisms. 2019;12:dmm036590.

37. White AB, Givogri MI, Lopez-Rosas A, Cao H, Van Breemen R, Thinakaran G and Bongarzone ER. Psychosine Accumulates in Membrane Microdomains in the Brain of Krabbe Patients, Disrupting the Raft Architecture. The Journal of Neuroscience. 2009;29:6068–6077.

38. Yamamoto K and Ando J. Endothelial cell and model membranes respond to shear stress by rapidly decreasing the order of their lipid phases. Journal of cell science. 2013;126:1227–1234.

39. Steinbuck MP, Arakcheeva K and Winandy S. Novel TCR-Mediated Mechanisms of Notch Activation and Signaling. The Journal of Immunology. 2018;200:997–1007.

40. Goenezen S, Rennie MY and Rugonyi S. Biomechanics of early cardiac development. Biomechanics and modeling in mechanobiology. 2012;11:1187–204.

41. Doubilet PM, Benson CB and Chow JS. Long-term prognosis of pregnancies complicated by slow embryonic heart rates in the early first trimester. Journal of Ultrasound in Medicine. 1999;18:537–541.

42. Braga M, Moleiro ML and Guedes-Martins L. Clinical Significance of Ductus Venosus Waveform as Generated by Pressure-volume Changes in the Fetal Heart. Curr Cardiol Rev. 2019;15:167–176.

