## Supplementary figures and images for "Mechanical forces pattern endocardial Notch activation via mTORC2-PKC pathway"

### Figure1-supplemnt figure S1

##### Figure 1-figure supplement 1

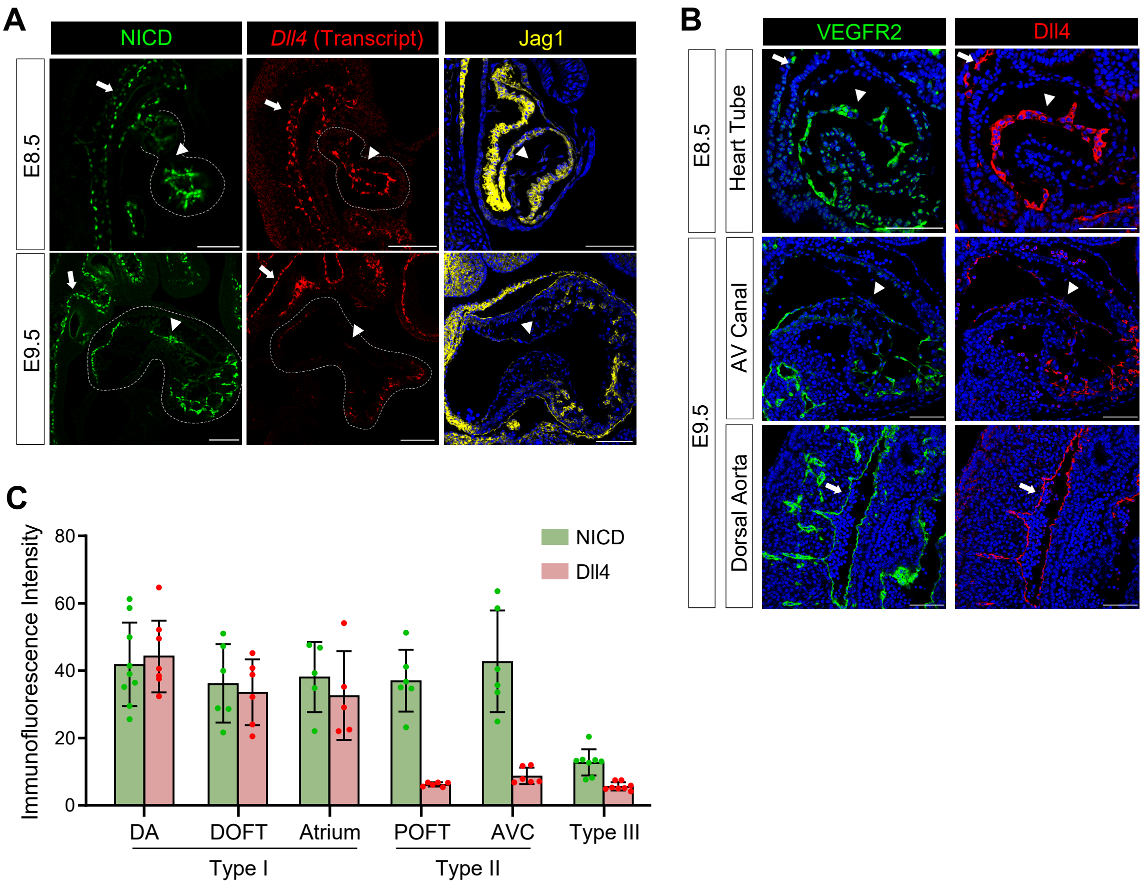

### Figure2-supplemnt figure S2

#### Figure 2-figure supplement 2

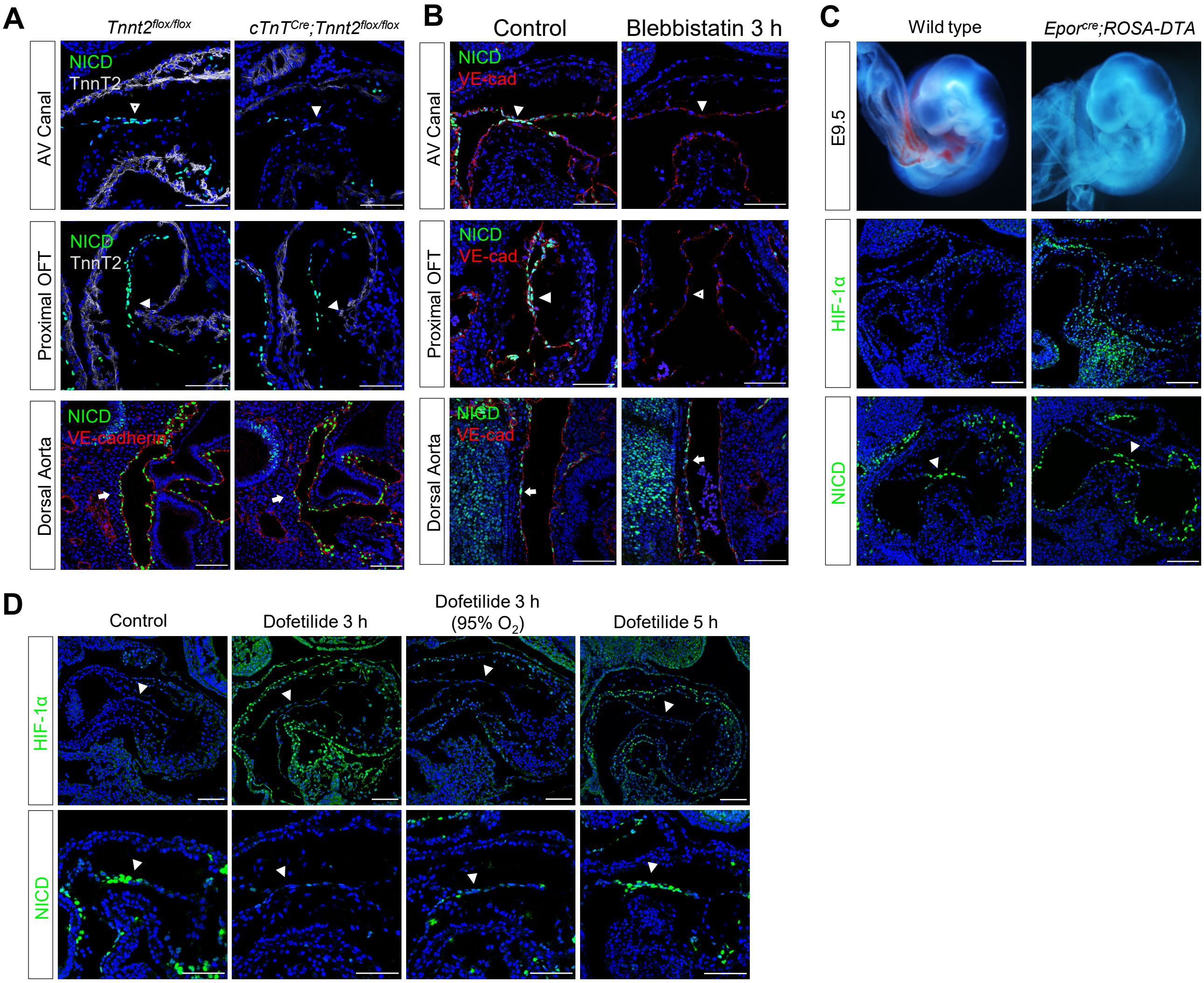

### Figure3-supplemnt figure S1

#### Figure 3-figure supplement 1

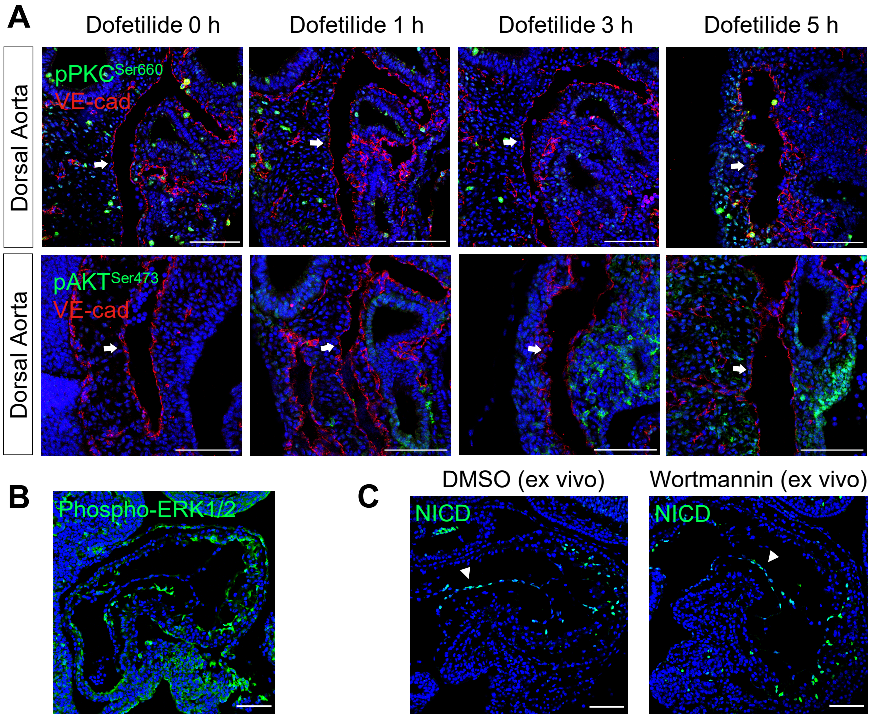

### Figure3-supplemnt figure S2

#### Figure 3-figure supplement 2

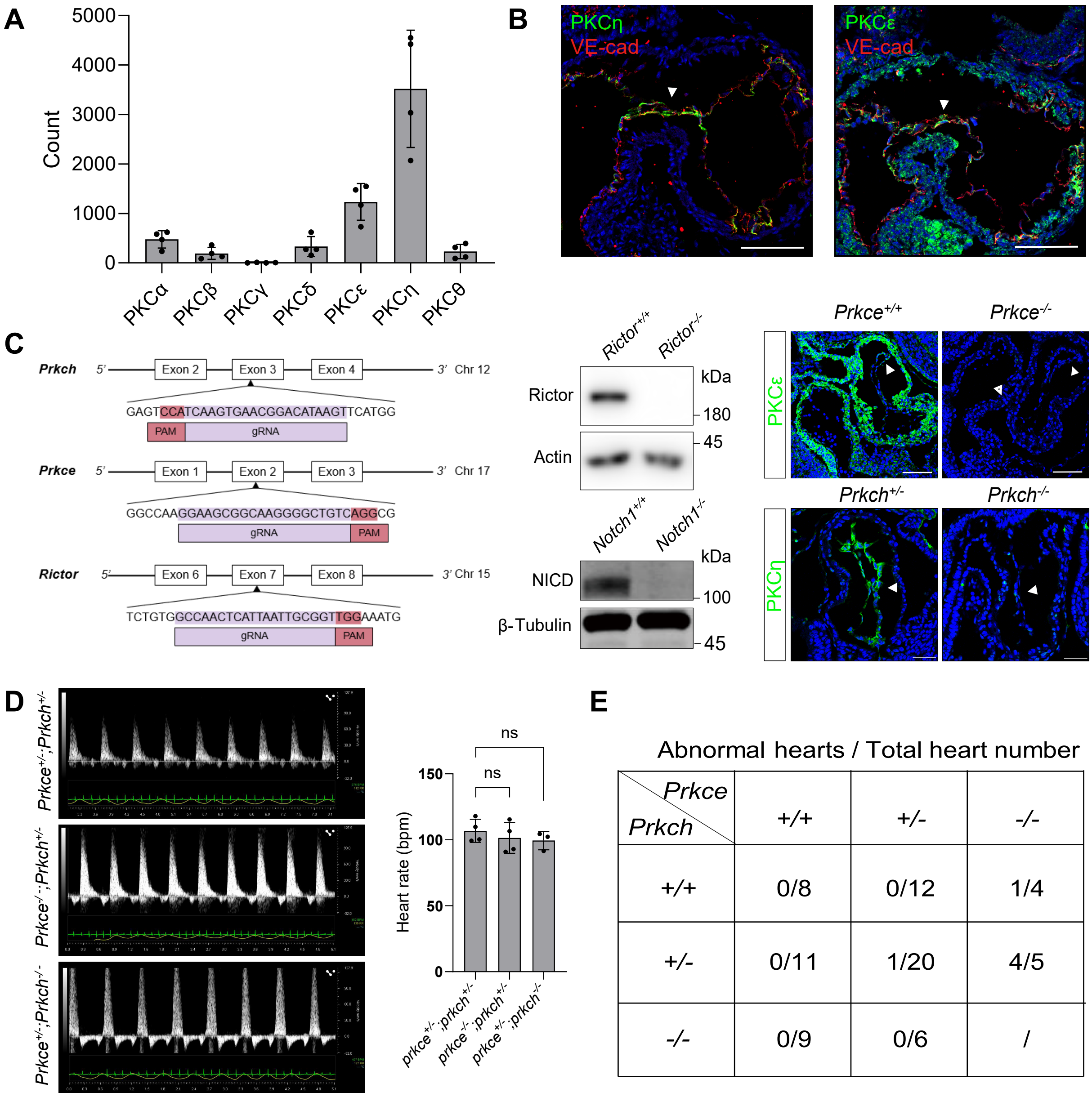


Abnormal hearts / Total heart number

| Prkch \ Prkce | +/+ | +/- | -/- |
|---|---|---|---|
| +/+ | 0/8 | 0/12 | 1/4 |
| +/- | 0/11 | 1/20 | 4/5 |
| -/- | 0/9 | 0/6 | / |

### Figure4-supplemnt figure S1

##### Figure 4-figure supplement 1

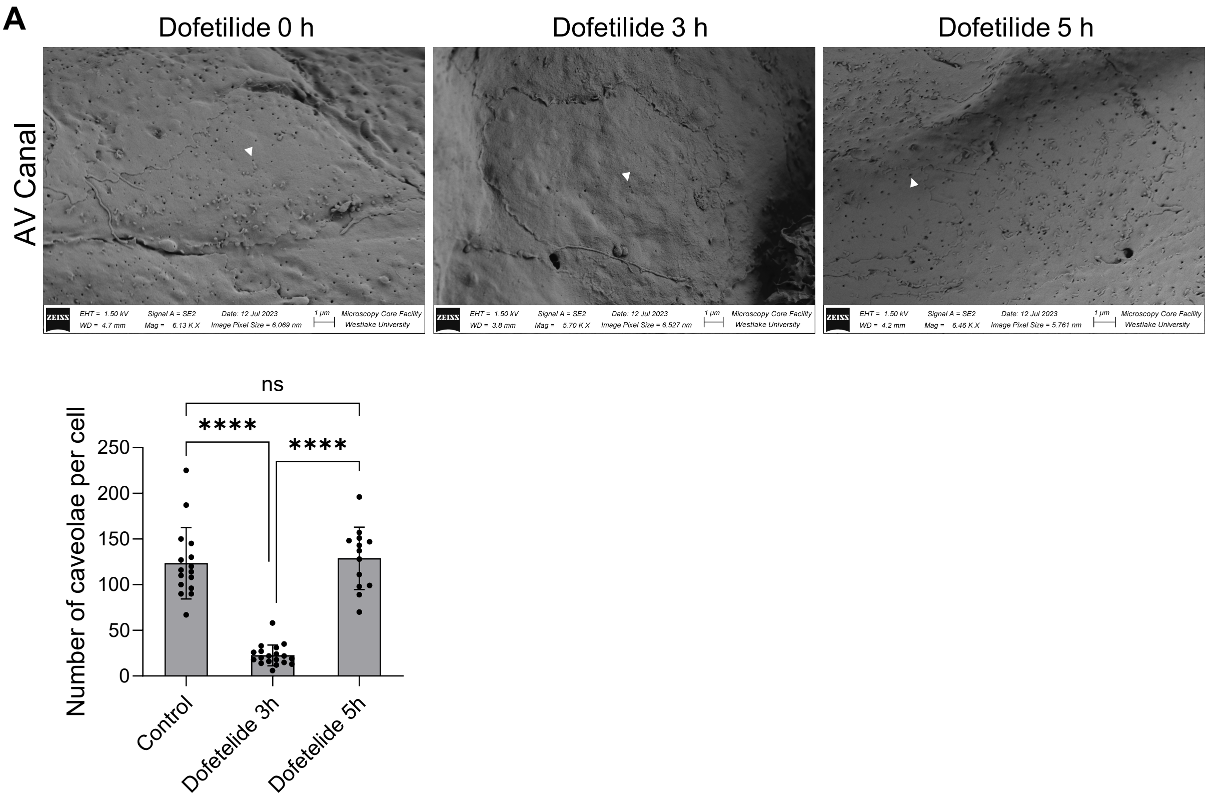
