## Supplementary material for "Mechanical forces pattern endocardial Notch activation via mTORC2-PKC pathway": Figure2-supplemnt figure S1

### Figure 2-figure supplement 1

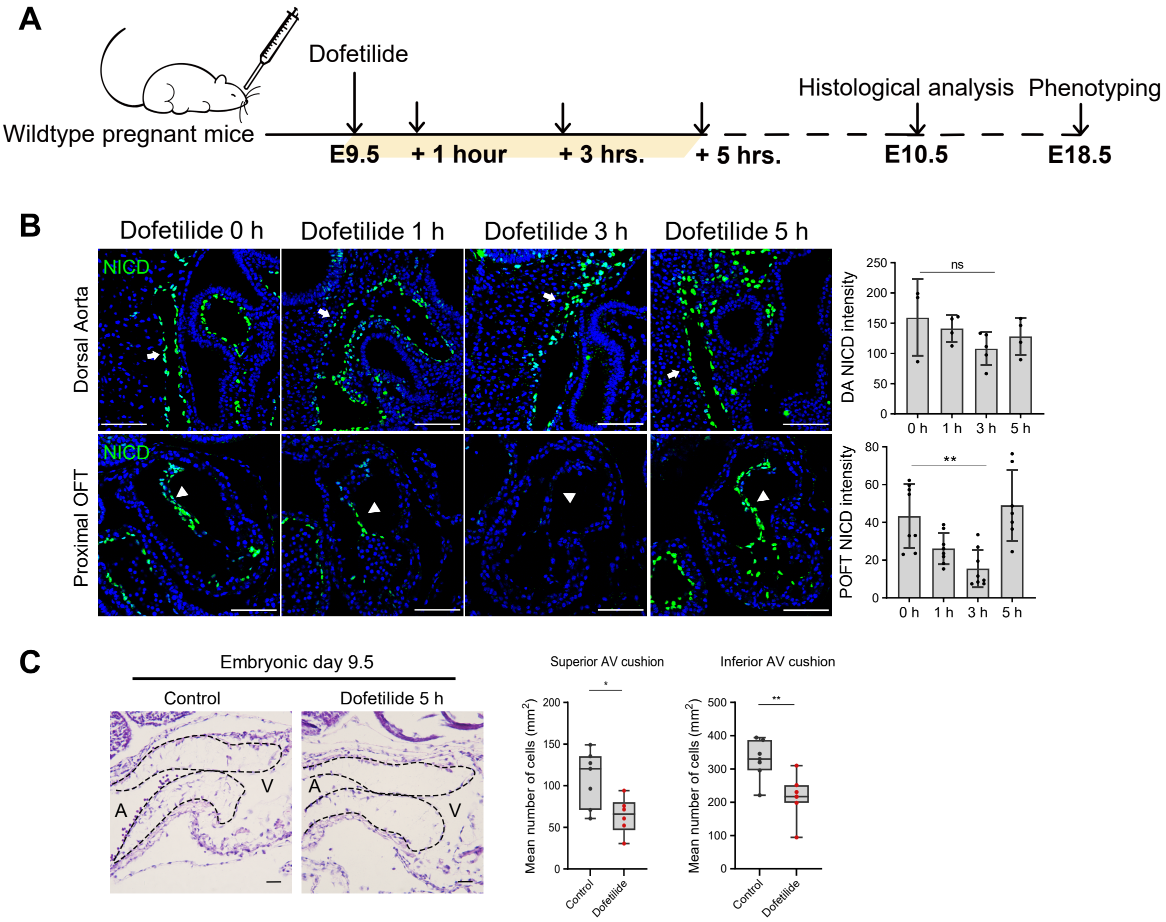


D

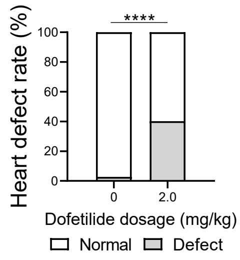


| Total No. of abnormal hearts | | 47 |
|---|---|---|
| Defect types | pmVSD | 30 (63.8%) |
| | DORV | 19 (40.4%) |
| | mVSD | 11 (23.4%) |
| | OA | 10 (20.3%) |
| | BAV | 8 (17.0%) |
| | AVSD | 6 (12.8%) |
| | BPV | 1 (2.1%) |
